# Determining cardiac structure by diffuse reflectance of different wavelengths

**DOI:** 10.1101/2021.09.23.460276

**Authors:** Philip Gemmell, Martin Bishop

## Abstract

Catheter ablation in patients suffering from chronic arrhythmias requires detailed knowledge of the underlying cardiac anatomy; such real-time, high resolution mapping is currently unavailable in a clinical setting. We present here preliminary work towards a novel optical strategy based on diffuse optical reflectance to provide quantitative anatomical measurements of the cardiac structure, including tissue thickness and presence of scar. An in-depth literature search is conducted to collate available experimental data regarding optical parameters in cardiac tissue and scar. Computational simulations of photon movement through cardiac tissue using Monte Carlo modelling are performed, with analysis being focussed on the effects on surface emission profiles of (i) optical parameters; (ii) tissue thickness; (iii) presence of scar. Our results demonstrate (i) sensitivity of the approach to changes in optical parameters within tissue, (ii) difference of results depending on light wavelength. These suggest that this can be used to detect cardiac anatomical structure to a depth of ∼ 2 mm, for both thickness of cardiac tissue and presence of scar. This study demonstrates the feasibility of using diffuse optical reflectance to determine cardiac structure, enabling a potential route for high-resolution, real-time structural information to guide catheter ablation and similar surgeries.

## 1 Introduction

Catheter ablation is a common clinical therapy to treat patients suffering from chronic arrhythmias [1], [2]. It involves the use of radio-frequency energy to destroy the electrical functioning of the targeted tissue, hence removing the conduction pathway that sustains the arrhythmia and terminating it, while also preventing re-occurrence. For this procedure, it is important to know the tissue properties: to know how much of the myocardium needs to be ablated, and where this ablation needs to be performed. If too little of the site is ablated, the underlying conditions for arrhythmia may remain, whereas if too much of the myocardium is sacrificed, a different host of problems arise, ranging from impaired cardiac contraction to perforation [3].

Current methodologies for imaging the cardiac structure are not ideally suited to this task. CT or MR scans do not provide the resolution sufficient to the task, and when such high resolution data do exist, it is often difficult to accurately map these pre-operative data to the real-time catheter location [1]. Alternative imaging modalities that can present high-resolution data in a real-time setting thus represent an important clinical goal.

Diffuse optical tomography presents a promising candidate. The use of optical tomography generally has a long history in clinical settings, using a wide variety of different approaches. Save for the eye [4], most biological tissues are relatively opaque, and thus imaging techniques must work to either reconstruct images using what ballistic photon data are available [5], or minimise the effect of scattering on reflected light, or else take account of it [6]–[8]. Optical coherence tomography looks to utilise the change in coherence due to differences in optical path length between ‘normal’ and tissue, imaging tissues to a high resolution [9]; this can be extended using fluorescence imaging [10]. However, this technique is limited in how deeply it can image, as it operates on the assumption of one (or very few) scattering events. Other techniques attempt to accommodate more, but still relatively few, scattering events, using confocal microscopy to select images based on two or more scattering events [11]. Other techniques, however, use data of the changes of optical properties between different tissue regimes to reconstruct the underlying structure based on how these changes influence the optical path [12]. Similarly, changes in the light transmission through tissue caused by changes in optical properties have been used to image to high resolution changes in hæmoglobin concentration by non-contact methods [13], [14].

So-called laminar optical tomography uses diffuse reflectance (potentially combined with confocal microscopy) to image structure to high resolution through limited tissue depths: the light undergoes many diffusion events during passage through the medium before being detected. This has been used extensively in neural applications [15], initially imaging vascular structure in the cortex due to absorption contrast [16], but by addition of fluorescent molecular imaging, its use has expanded to structural analysis [17], [18]. However, its investigation for use in guiding catheter ablation is still in its infancy, with previous work being preliminary in its thoroughness of parameter effects [19]. This paper looks to address this by providing a comprehensive literature review for cardiac optical properties, and using the data to comprehensively assess the potential for optical tomography to determine cardiac structure.

The imaging modality presented herein relies on diffuse optical reflectance, wherein light is shone onto a surface, before being scattered and diffused through the medium before being re-emitted and detected at the surface of illumination. Light which exits the tissue at a distance further from the initial source location will have, on average, penetrated to a greater depth within the tissue. By suitable analysis of the surface emission profile, it is thus hoped to be able to extract data regarding the tissue structure. To conduct accurate simulations of this phenomenon, it is necessary to establish the optical properties of cardiac tissue, both normal and scar. There is a wealth of experimental data in the literature for optical properties of different types of tissue; however, these data are diffuse, and there does not exist a comprehensive review of the values that can be found.

In this work, analysis will be performed to determine what data can be extracted from the diffusely reflected light: whether the emission profile, and changes therein, can be usefully analysed to reveal details regarding tissue structure (both thickness and presence of scar). It is assumed that thinner tissue will curtail the deeper penetrating banana paths, in turn affecting the emission profile, while detecting the presence of sub-surface scar is based on (assumed) changes in optical properties of the tissue (such as *µ*_*a*_ and *µ*_*s*_) affecting the photon paths. These optical parameters can be poorly defined, with conflicting values given in the literature (see Supplementary Information); differences in these parameters between normal and scar tissue are little reported. To that end, a comprehensive literature search is undertaken to collate the values given for various optical parameters. Following that, the effect of changes in optical parameters are investigated to establish their relative effects and influences on the emission profiles. The emission profile is then examined using various different parameter values to judge the efficacy of optical tomography in establishing (i) tissue thickness and (ii) presence of scar tissue.

## 2 Materials & Methods

### 2.1 Theory & Literature Search

The path taken by individual photons in this scattering/diffusion process is stochastic and thus cannot be predicted, but through the use of large-scale computer simulations it is possible to assess the average properties of the photon path. These average properties will reflect data regarding the depth of the photon penetration into tissue, with those being re-emitted at the surface having been diffusely scattered extensively for their trajectory to return to the surface. This is referred to as the ‘banana effect’ due to the shape of the most common photon paths. These ‘banana paths’, and the resulting surface emission profiles, depend on the optical properties of the medium through which the photons travel. Changes in the emission profiles can be traced back to changes in the banana paths of the photons, which in turn depend on the tissue, and the optical properties of that tissue. The importance of optical properties as an imaging modality has already been established: the changes in optical properties, and their influence on light diffusion, have been used to detect changes in hæmoglobin, though not the structure therein [13]. It is expected that the optical properties vary depending on not only tissue structure (normal tissue *versus* scar tissue, for example), but also on the wavelength of the light used: different wavelengths of light (with longer wavelength light typically penetrating further into tissue) will add further detail to the picture.

Determining the optical properties for tissue is a non-trivial process, and is subject to many complicating factors. For example, changes in hæmoglobin, tissue water content and (significantly for this work) light wavelength are known to have noticeable effects on optical properties [20]. As part of this work, an extensive literature search was conducted for values of optical properties of tissue recorded in the literature. This was conducted through a combination of tracing the literature through other recent works to find the experimental root, examining those works that cite seminal works (*e*.*g*. [21]), and PubMed searches for terms such as ‘light tomography’.

Firstly, it is necessary to confirm that changes in the optical properties of bulk tissue (and hence potentially the presence of scar with different optical properties) can be detected by changes in the emission profile. As such, simulations were done to assess the effects of changes in 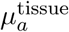 and 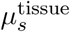 on the emission profiles from tissue. Subsequently, the effect of tissue thickness was modelled, followed by the presence and location of scar tissue.

### 2.2 Model Set-Up

In determining the validity of using diffuse reflectance of light to determine the structural properties of cardiac tissue, computational simulations of the diffusion, reflectance, absorption and transmission of light through cardiac tissue must be completed. In constructing this experimental set-up, we simulate a point source beam of light incident at the origin of a flat plane of cardiac tissue. The path of individual photons is simulated through Monte Carlo modelling (described in detail in the following section). While most photons simulated travel through the tissue and extinguish within the tissue, some, via diffusion and reflection, are re-emitted at the surface of the tissue—this is shown in Fig. 1. The ‘weight’ of the photons that are re-emitted (expressed as a percentage of the power of the initially injected photon) are recorded at their radial distance from the point of injection.

**Figure 1:**
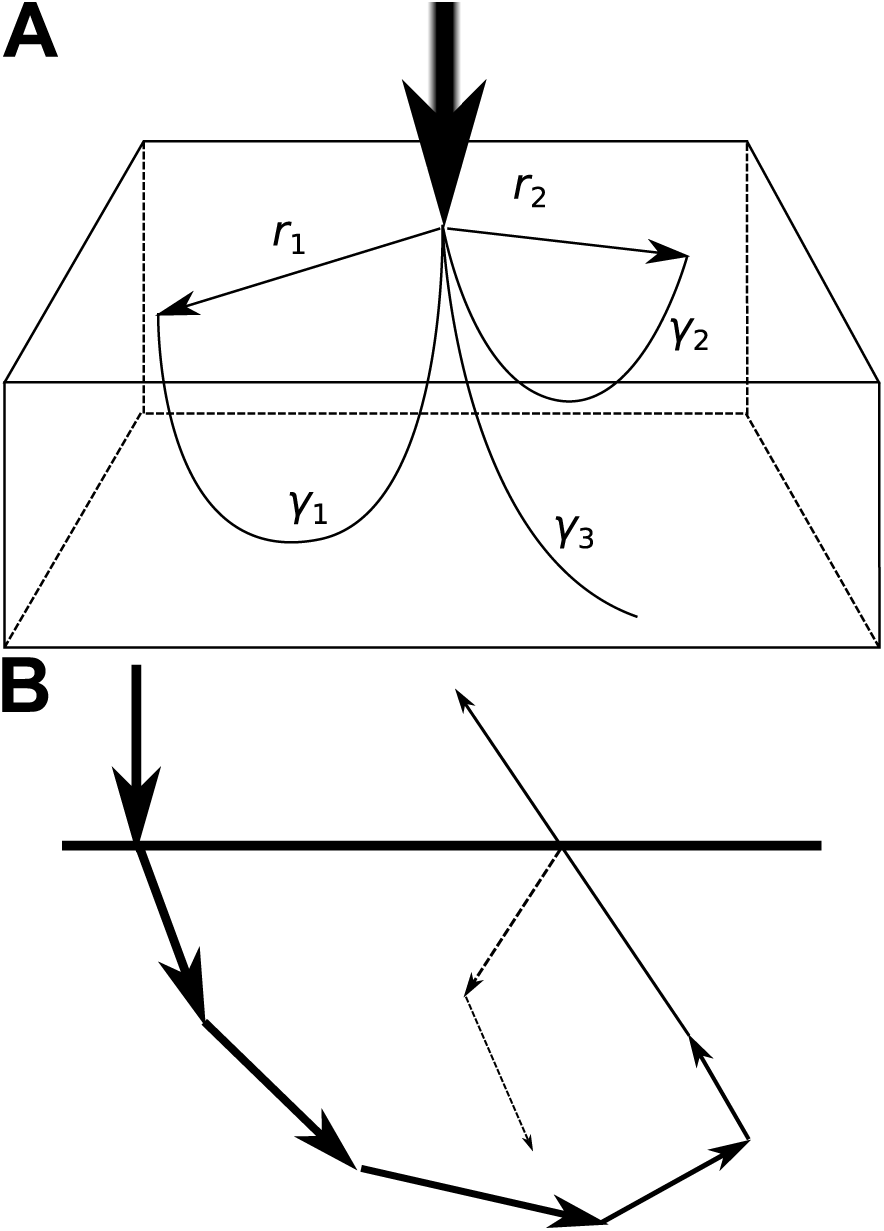
Schematic demonstrations of the photon trajectories being simulated. (A) Experimental set-up to study diffuse optical tomography. Three example photon traces are shown, two of which (γ_1_ and γ_2_) represent reemission at two different radii, r_1_ and r_2_. A third photon, γ_3_, is not re-emitted, and is extinguished within the tissue. (B) Schematic example of a photon’s path through tissue. With each step, some absorption of the photon’s energy takes place, thus reducing the photon weight for the subsequent step. If the photon encounters the tissue boundary, it may either be transmitted (solid line) or reflected within the tissue (dashed line).

Most initial results are determined from simulations conducted for semi-infinite tissue, *i*.*e*. infinite in *x* and *y* directions, and extending to infinity from the zero-plane in *z*. Some simulations are conducted in tissue that has a finite extent in the *z* direction. Photons are injected into the tissue perpendicular to the *xy*-plane at the origin. Simulations and analysis are performed using Python3; simulations are performed using multi-core processing on a 24×2.5 GHz CPU.

### 2.3 Photon movement

Photon propagation through cardiac tissue is simulated using a step-by-step Monte Carlo modelling method, originally demonstrated in [21]. A brief summary of the methodology is presented here; for further details, the reader is referred to the original paper. Throughout the following, there are several references to *ξ*_*i*_: these are uniformly distributed random variables between 0 and 1.

The over-arching methodology is to simulate the movement of the photon using discrete stochastic step-sizes through tissue, with absorption and transmission being calculated at each step (Fig. 1B). The initial input into tissue assesses a photon packet of initial normalised weight 1; here, the photon beam is modelled to enter the tissue in the direction normal to the tissue surface. Initial specular reflectance reduces the photon weight as *W*_photon_ = 1 − *R*_*sp*_, where *R*_*sp*_ = (*n*_*t*_ − *n*_*m*_)^2^*/*(*n*_*t*_ + *n*_*m*_)^2^ with *n*_*t*_ representing the refractive index of the tissue and *n*_*m*_ the refractive index of the outside medium. *n*_*m*_ is set to 1.0 (the refractive index of air), and *n*_*t*_ is set to 1.4 (as used in [19], [22], and representing a reasonable average of experimentally measured values (see S4 Table)).

Within the tissue, the photon is advanced by a step-size *s* at each step of its journey. This step-size is sampled from a distribution (wherein 0 ≤ *s* ≤ ∞) according to *s* = − ln(*ξ*_1_)*/µ*_*a*_ + *µ*_*s*_, where *µ*_*a*_ and *µ*_*s*_ refer to the absorption and scattering coefficients respectively. The photon’s position is then updated from **x** to **x**′ according to 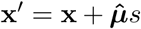, where 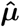 is the unit vector describing the photon’s direction, and can be devolved to its component parts in the *x, y* and *z* directions as shown in Fig. 2 and calculated thus:

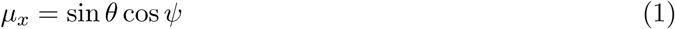

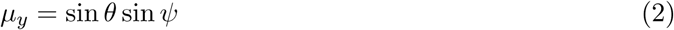

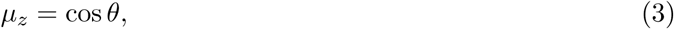

**Figure 2:**
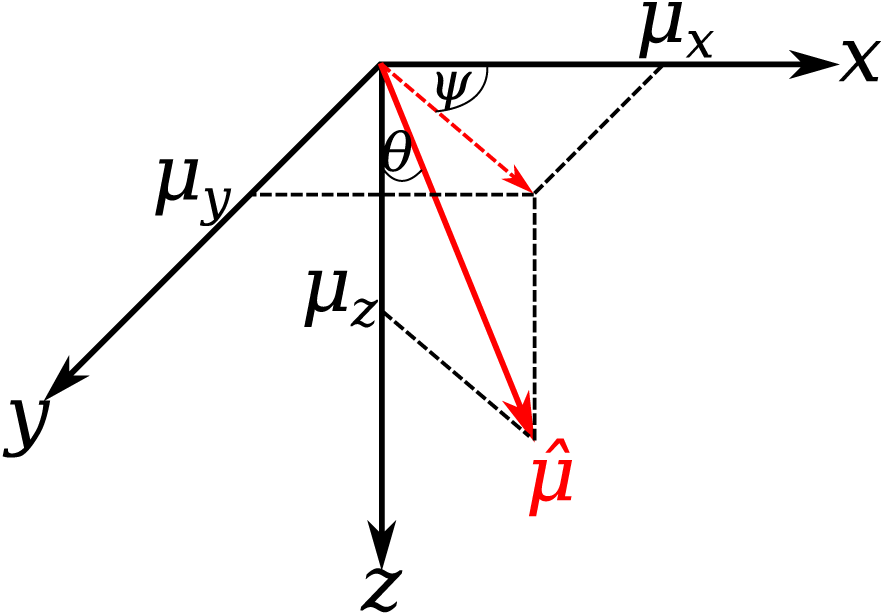
Angles and components of the direction of photon propagation.

where *θ* is the deflection angle from the *z*-axis, and *ψ* is the azimuthal angle made by the projection of 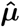 onto the *xy*-plane and the *x*-axis.

Following each step, photon absorption is modelled such that the packet deposits a proportion of its weight in the tissue at its current location. The amount deposited is given by 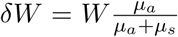, with the photon packet’s weight being correspondingly reduced by *δW*.

The photon’s direction is then updated, depending in part on the value of the tissue scattering coefficient *g*, which dictates the extent of forward- or back-scattering and thus the value of *θ*:

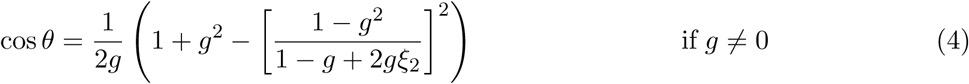

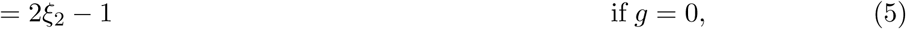

The azimuthal angle does not have this dependence on *g*, instead being uniformly distributed between 0 and 2*π* (*ψ* = 2*πξ*_3_). The components of 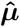 are then updated to 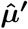 accordingly:

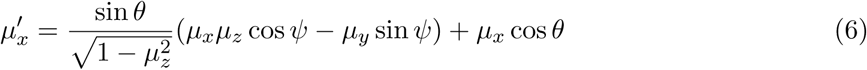

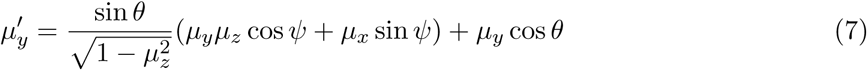

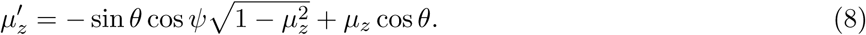

However, if the photon direction is sufficiently close to the *z*-axis (*µ*_*z*_ > 0.99999), the direction is instead updated as

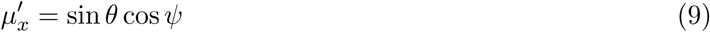

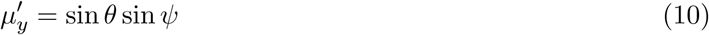

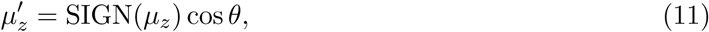

where SIGN(*µ*_*z*_) returns 1 when *µ*_*z*_ ≥ 0, and −1 otherwise.

The photon packet’s journey ends under one of two conditions: termination or transmission. In the former case, when the packet’s weight falls below a threshold value (set as *W*_threshold_ = 0.0001), there is little to be gained from further simulation. However, to ensure conservation of energy and that the photon weight distribution is not skewed, the photon packet is terminated via Russian Roulette: the packet has a probability of 1*/m* of surviving with an updated weight of *Wm*, or else being permanently terminated, with *m* is set to a value of 10 [21], [23].

Alternatively, during the random walk of photons through the tissue, some photons will eventually encounter a tissue/medium boundary (Fig. 1). When a photon-tissue boundary interaction occurs, the photon is either internally reflected back into the tissue where its journey continues, or it is transmitted out of the tissue and no further simulation of its path is performed. If the photon interacts with the boundary with an angle of incidence *α*_*i*_ greater than the critical angle (*α*_*crit*_ = arcsin(*n*_*m*_*/n*_*t*_)), the photon is necessarily internally reflected. Alternatively, the reflection coefficient *R* is calculated using Fresnel’s formulae:

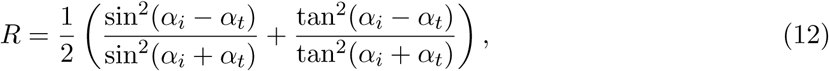

where *α*_*t*_ is the angle of transmission (calculated using Snell’s Law, *n*_*m*_ sin *α*_*t*_ = *n*_*t*_ sin *α*_*i*_). If *R < ξ*_3_, then the photon is transmitted; otherwise, the photon is internally reflected, and the photon position updated accordingly. If a photon is transmitted at the tissue surface, *i*.*e*. on the same place into which the photon was originally injected, the weight of the photon and radial distance from the site of injection is recorded for analysis; transmission at any other boundary is recorded as termination, for our purposes.

### 2.4 Validation of model and normalisation

The absorbed photon density within tissue can be approximated by a monoexponential decay function, Φ = Φ_0_ exp^*−r/δ*^, with the photon density Φ decaying as it travels a distance *r* through the tissue, with the 1*/e* attenuation depth given by *δ*. This penetration depth can be estimated (*δ*_eff_, Fig. 3A), and compared with that value predicted by the analytic solution to the photon diffusion equation for a point source over a geometrically regular domain 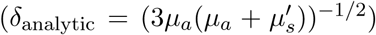 [24]. For the example shown in Fig. 3A (*µ*_*a*_ = 0.23 mm^-1^, *µ*_*s*_ = 15.0 mm^-1^), *δ*_eff_ = 1.1990, with *δ*_analytic_ = 1.1325.

**Figure 3:**
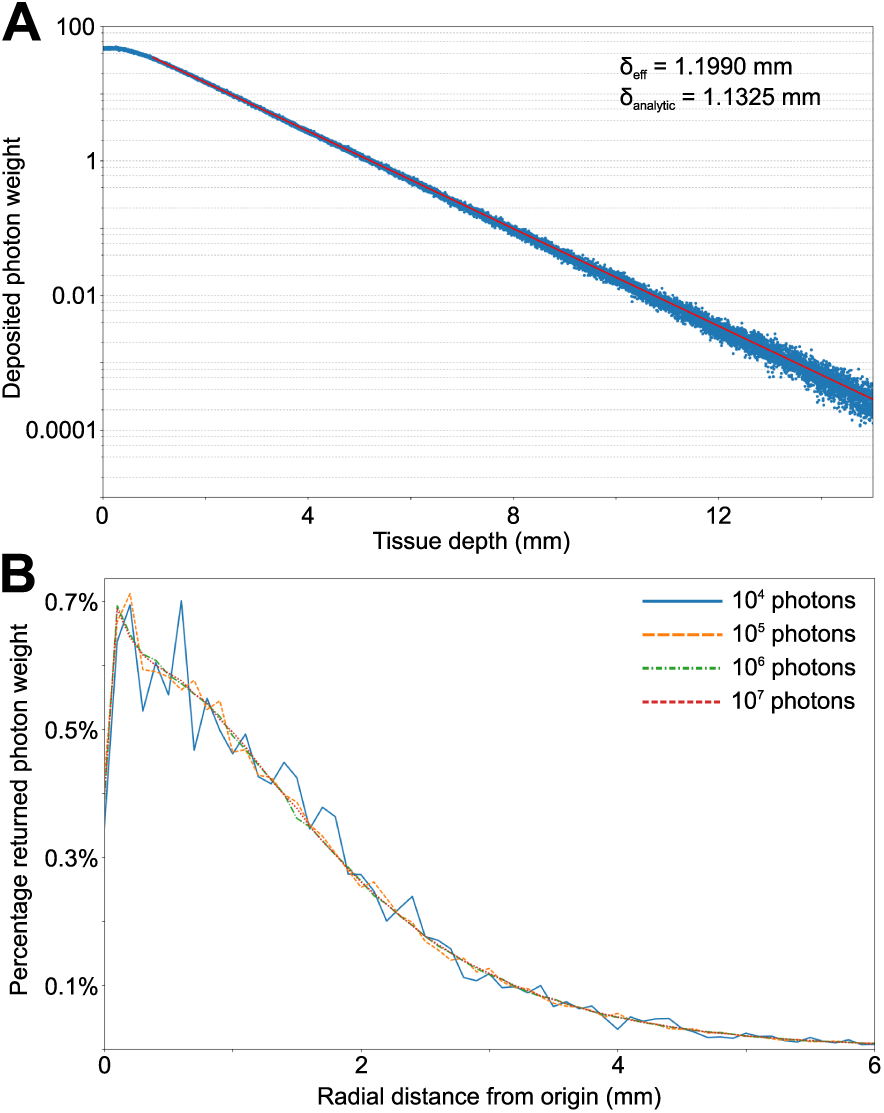
Validation of the photon simulation method. (A) Absorption of light (10^5^ photons) in infinite tissue, with the line of best fit for the derivation of δ_eff_ shown in red. (B) Comparison between normalised surface emissions from infinite tissue.

It was also established that the results of simulations are, save for stochastic effects, independent of the number of photon packets simulated. This is shown in Fig. 3B, which compares surface photon emission traces normalised to the incident photon weight. No simulation was performed for fewer than 10^4^ photons, that number being the minimum scale of Monte Carlo simulations to achieve statistical significance [25].

### 2.5 Data Analysis

All photons that are transmitted at the tissue surface have their residual weight recorded (*i*.*e*. the relative weight they possess compared to their initial weight at the start of their journey), and their radial distance from the origin at their point of emission. This permits analysis of the relation between the emitted photon weight *vs*. the radial distance from the site of photon injection—an example trace is shown in Fig. 4, which also demonstrates the terminology used in this paper. The emission profile directly correlates to the probability distribution for the radial distance from origin that photons will travel before returning to the tissue surface (it is not related to the probability distribution for all photons’ travel distances).

**Figure 4:**
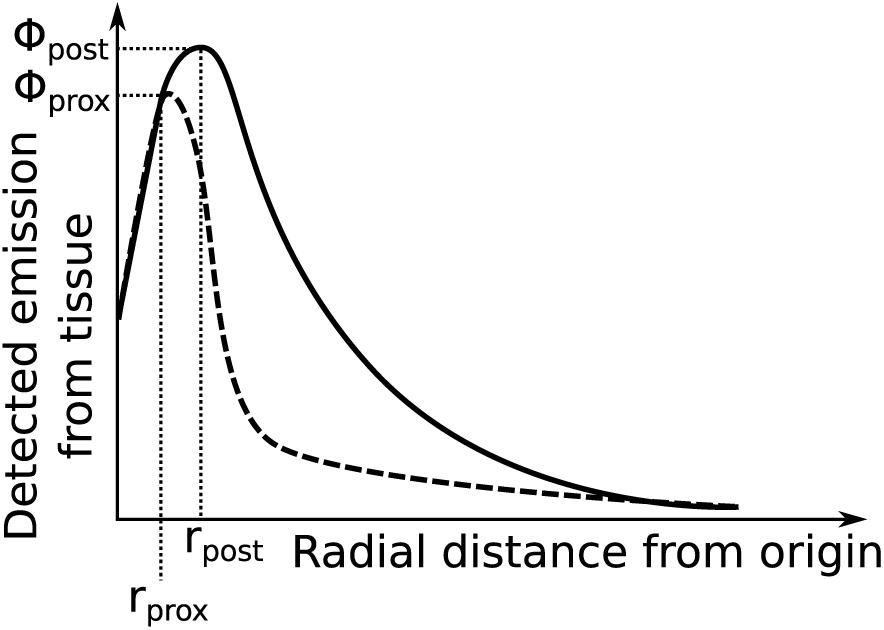
Schematic of expected photon emission from tissue Φ as a function of radial distance from site of photon injection. r_prox_ and Φ_prox_ refer to the radius and photon emission strength respectively immediately proximal to the source, while r_post_ and Φ_post_ refer to the radius and photon emission strength respectively for a subsequent peak. It should be noted that Φ may not increase beyond Φ_prox_, and thus r_post_ and Φ_post_ would be undefined (dashed emission profile).

There will be a non-zero element of photon emission for *r* = 0, reflecting those photons whose path leads them to return to the origin. Under most circumstances, however, it is expected that the probable photon path will lead to re-emission at some radial distance from the origin; depending on the optical properties of the tissue, and thus the photon path through tissue, the peak photon emission (Φ_peak_) will be at *r* > 0.

Changes in the emission profile can be measured: for example, changes in Φ_peak_ and the full-width half-maximum (FWHM) of the emission profile. It is hoped that there are correlations between changes in the emission profile changes and various alterations in the tissue (tissue thickness, presence of scar, etc.). These changes may be revealed either directly, or by the differential change between emission profiles observed for light of different wavelengths.

## 3 Results

### 3.1 Effects of 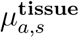

The baseline parameters used for 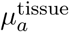 and 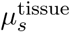 simulations were *µ*_*a*,0_ = 0.3 mm^-1^ and *µ*_*s*,0_ = 15.0 mm^-1^, which represent the approximate average values observed across a range of wavelengths for cardiac tissue (see Supporting Information). Simulations were conducted for 10^5^ photons injected into semi-infinite tissue; results are shown in Fig. 5. The left panels of this figure plot the maximum depth each photon packet reached before returning to the surface for emission, *versus* the radial distance from the site of entry for that emission; these confirm the working hypothesis that those photons that are re-emitted at a greater radius have, on average, penetrated to a greater depth within the tissue. The right panels show the corresponding radial emission profile.

**Figure 5:**
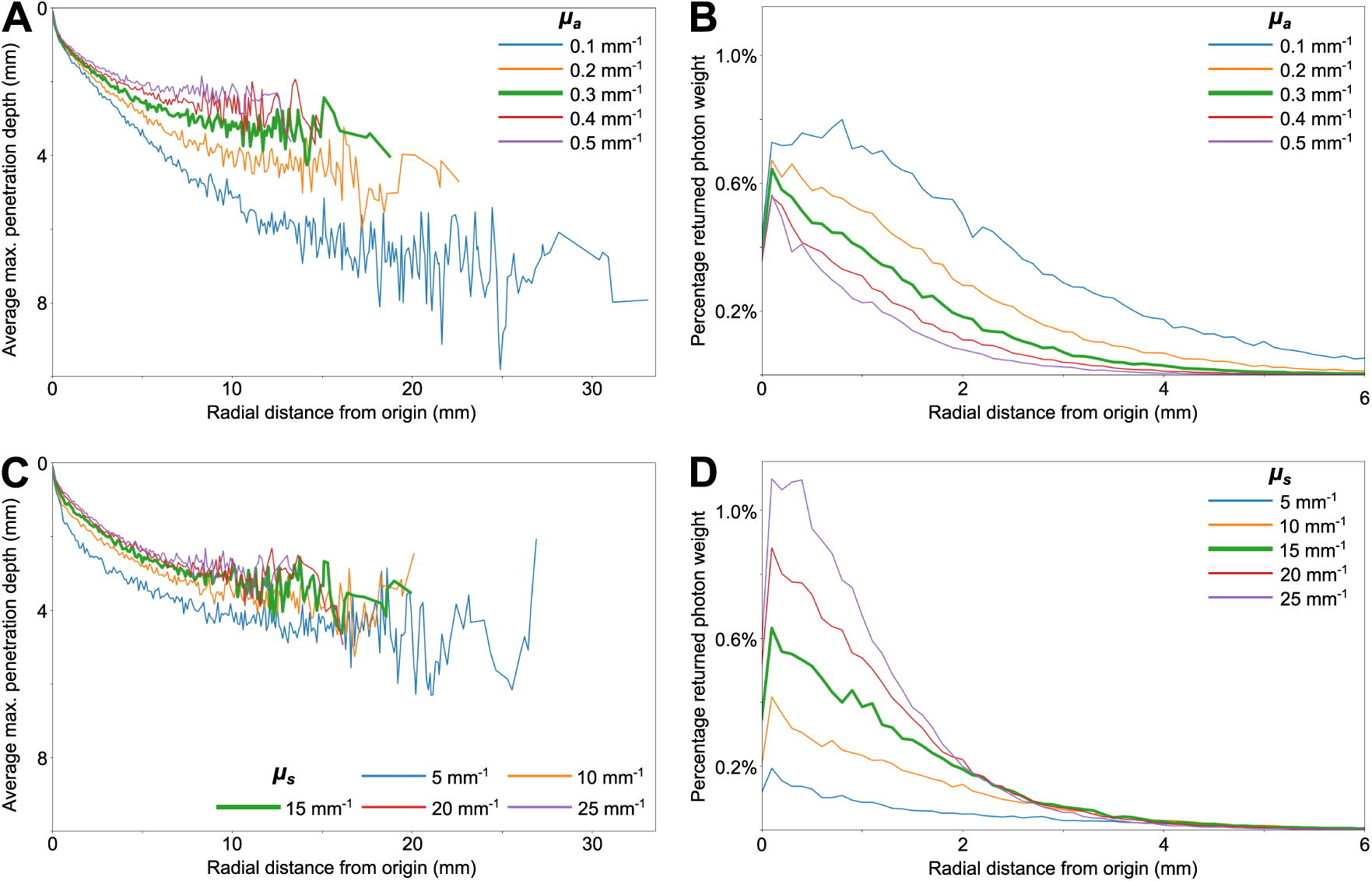
Changing properties of photon emissions for changes in 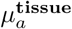 and 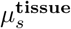. The baseline parameters are shown in each plot by the bold green line. (A,B) represent data for changing µ_a_, (C,D) represent data for changing µ_s_. (A,C) plot tissue penetration depth v. radius of surface emission. (B,D) plot photon emission weight v. radius of surface emission.

Variation of 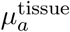 leads to noticeable changes in the photon penetration through tissue, which is in turn detectable via the emission profile at the tissue surface. A reduction in 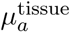 leads to an increase in penetration depth for a given radius of detection, *i*.*e*. when photons are detected upon emission at the tissue surface, they have (on average) penetrated to a greater depth within the tissue (Fig. 5A). For example, for 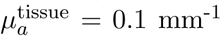 those photons emitted at *r* ≈ 10 mm have penetrated to almost 5.5 mm within the tissue, whereas for 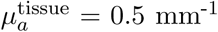 the photons have, on average, only penetrated 2.5 mm within the tissue. Not only does this decrease in 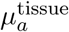 lead to greater tissue penetration for a given radial emission, but the maximum radial range is increased: data for 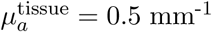 extends to *r* ≈ 13 mm, whereas 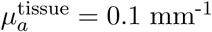 records emissions for *r* ≈ 33 mm. However, examination of the surface emission profile reveals that the photon emission at these radii may well be too small to be detectable regardless.

There are distinct changes in the surface emission profile: Φ_prox_ increases as 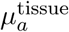 decreases, but, more significantly, the rate of decline in Φ after *r*_prox_ decreases (Fig. 5B). This is reflected by an increase in the full-width half maximum (FWHM) of the emission profile as 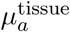 is decreased for all values of 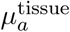. For small values of 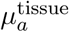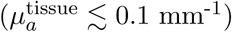, the change in Φ_post_ is such that it exceeds Φ_prox_.

Changes in 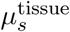 produce notably different effects, in both the absorption and emission profiles. As 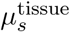 increases, the depth of penetration in tissue for photons emitted at a given radius is reduced (Fig. 5C). This effect is more noticeable within a shorter radius of emission than changes in 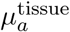 (compare Fig. 5C with Fig. 5A for small radii of emission). However, differences in 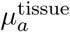 have a greater effect at greater radii.

This difference in tissue absorption at small radii is reflected in the surface emission profile (Fig. 5D), whereby an increase in 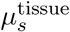 results in an increase in Φ_prox_. However, unlike with 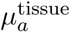, the decay in Φ from Φ_prox_ does not alter significantly: changes in 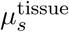 appear to be more readily observable via the surface emission profile at small radii, with these changes becoming negligible for *r ≳* 4 mm. Beyond this radius, changes in 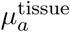 would thus appear to be the primary cause.

‘Average’ parameters are based on those values found in the literature search, reported in the Supporting Information.

### 3.2 Effect of tissue thickness

Simulations were then repeated on tissue slices of varying thickness, using both the average parameters defined in the previous sections, and the parameters noted in [26] and [27] (referred to as Ding and Walton parameters, respectively). While the averaged parameters are indicative of overall behaviour, these papers and the parameters therein (Table 1) are significant in allowing examination of the effects on different wavelengths, with the associated parameters determined under consistent experimental conditions; under a clinical setting, the *relative* changes in emission profile properties for different wavelengths would be the relevant variable, as the absolute properties will be uncalibrated.

**Table 1:** Optical parameters used in simulations.

| Source | Light wavelength (nm) | $\mu_a^{\text{tissue}}$ ($\text{mm}^{-1}$) | $\mu_s^{\text{tissue}}$ ($\text{mm}^{-1}$) |
| --- | --- | --- | --- |
| ‘Average’ | n/a | 0.23 | 15.00 |
| [27] | 532 | 0.88 | 7.42 |
|  | 715 | 0.05 | 5.40 |
| [26] | 488 | 0.52 | 23.00 |
|  | 669 | 0.10 | 21.80 |

Fig. 6 plots the radial emission profiles as a function of tissue thickness, using average optical properties. For tissue thicknesses (*t*) greater than ∼ 2 mm, there is negligible difference in observed emission profile. For *t* = 1mm, there is a notable increase in Φ(*r* ≈ 1mm), likely due to the photons that are reflected at the bottom surface of the tissue reaching the surface of the tissue to affect the emission profile. As the tissue thickness decreases further, the radius at which this reflection effect is observed reduces, due to the photons having less distance to scatter radially before being reflected and emitted. At the same time, the increase in Φ due to these reflected photons increases, due to a greater proportion of photons being reflected and emitted prior to being extinguished due to the shorter travel time through tissue.

**Figure 6:**
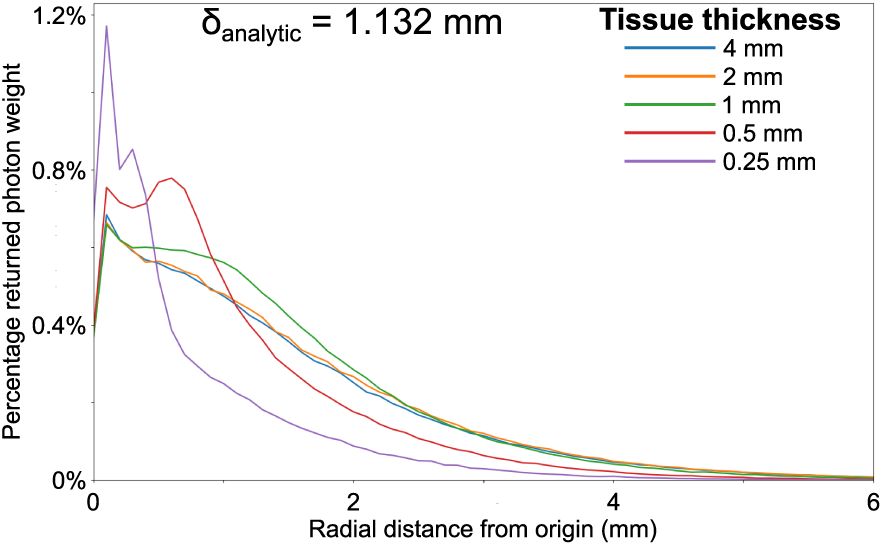
Effect of varying tissue thickness. Photon emission profile for tissue of varying thickness using average optical properties 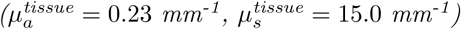.

Fig. 7 show the surface emission profiles for different tissue depths, using parameters appropriate for Walton (top panels) and Ding (bottom panels). It can be seen that the findings for the ‘average’ parameters are broadly reproduced, though there remain substantial differences in the details. For both wavelengths of the Walton parameters, Φ_prox_ is dependent on tissue thickness: for 715 nm, differences can be noted for *t* ≲ 2 mm (Fig. 7B), whereas this is reduced to 1 mm for 532 nm light (Fig. 7A). Furthermore, the increase in Φ_post_ evident for average parameters is also evident for both 532 nm and 715 nm light: see Φ(*t* = 0.5mm) for 532 nm light, and Φ(*t* = 0.5mm, 1mm) for 715 nm light.

**Figure 7:**
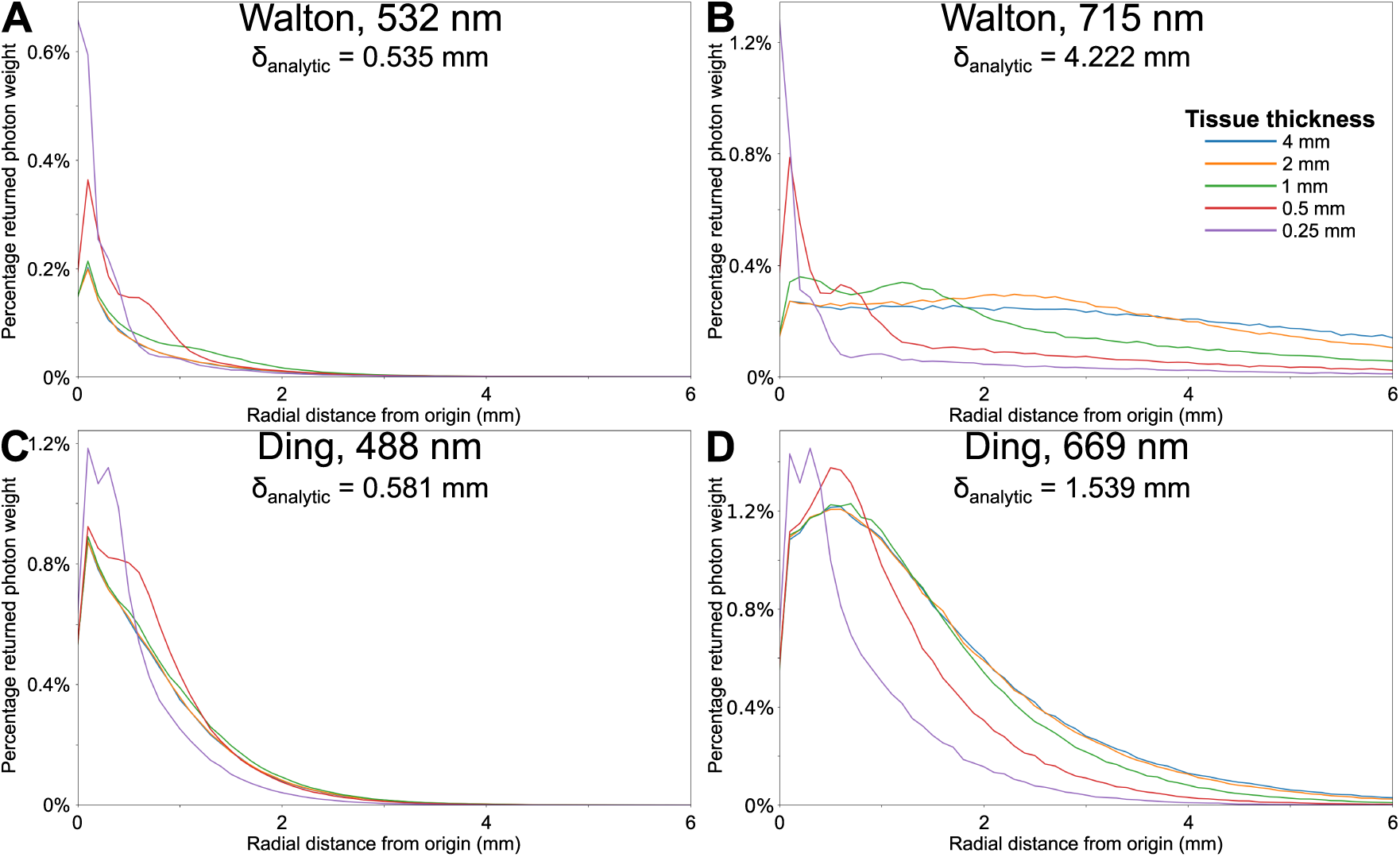
Effect of tissue thickness on surface emission profiles. Simulated using parameters from [27] and [26].

The difference for 715 nm data is due to the more penetrating nature of the light, wherein photon absorption in infinite tissue can still be detected in tissue ∼ 60 mm thick (data not shown). This increase in photon penetration also leads to noticeable differences in emission profiles for 715 nm at greater *r*, whereas for 532 nm light there is negligible difference in emission profiles for *r ≳* 2 mm.

The emission profiles derived from the Ding simulations demonstrate similar trends to the Walton simulations, if to a more muted degree. There is a less significant change in Φ_prox_ with tissue thickness; it can be noted that the simulations for 669 nm light demonstrated tissue penetration depths of ∼ 30 mm, which, while significant, is still less than for Walton 715 nm simulations. However, this relative consistency in Φ_prox_ makes subsequent changes in Φ(*r*) more easily detectable by changes in ratios. For example, change in FWHM for different tissue thicknesses are more readily apparent, especially for 669 nm.

### 3.3 Effects of scar depth

As changes in *µ*_*a*_ and *µ*_*s*_ can have significant impacts on the emission profiles, it is reasonable to expect that changes in these parameters between scar and tissue may have an equally profound effect. Optical properties of scar tissue are poorly reported in the literature [20], [28], and those values that are reported in the literature are not entirely in agreement. It is relatively consistently indicated that 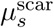 is reduced compared to 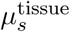, but the effect of scar on *µ*_*a*_ is less clear.

Fig. 8 shows the changing emission profiles for a scar 0.5 mm beneath the tissue surface in semi-infinite tissue with ‘average’ optical properties, with varying values assigned to 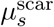 (Fig. 8A) and 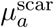 (Fig. 8B). No scar was simulated closer to the surface due to a surviving endocardial layer due to the perfusion of nutrients from the ventricular blood pool.

**Figure 8:**
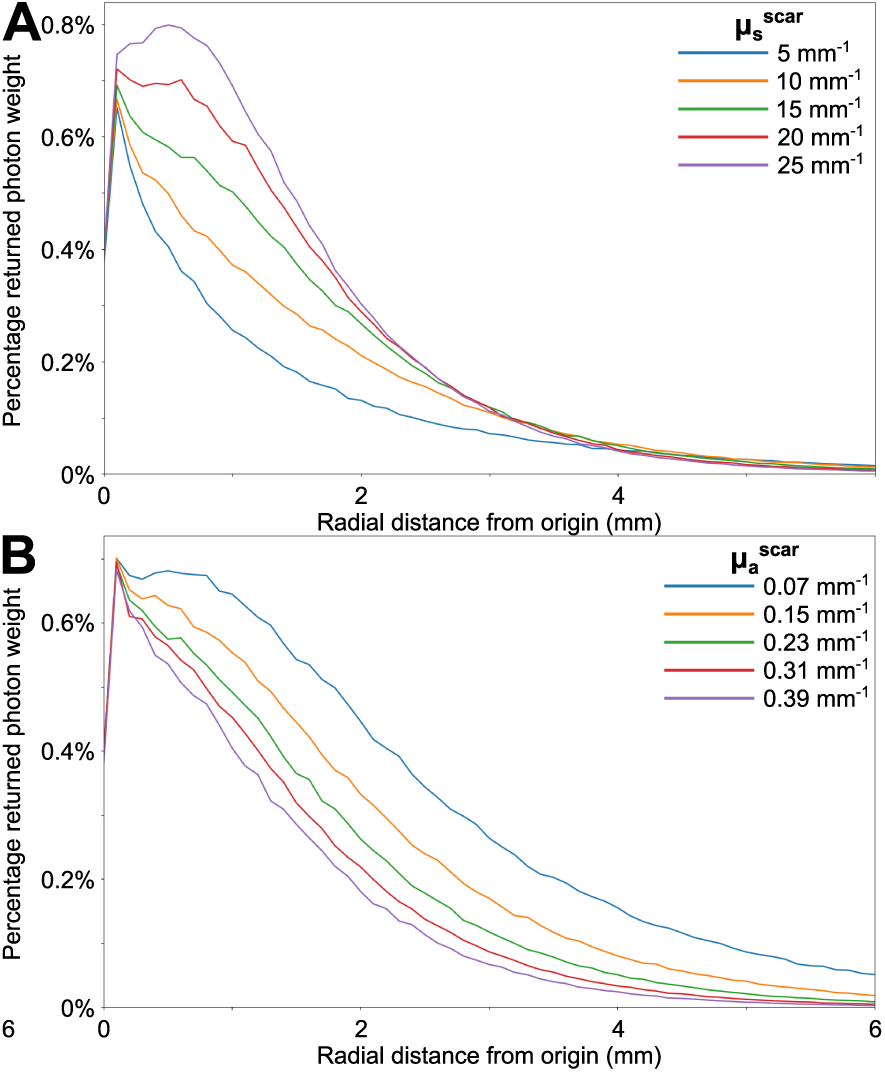
Effect of changing scar properties on surface emission profiles. Effect of scar 0.5 mm beneath the surface on photon emission from the surface of the tissue, with varying values used for (A) 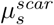 and (B) 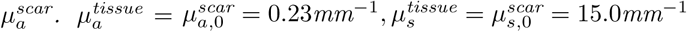.

An increase in 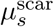 results in a reduced rate of decay of photon emission with radial distance. For large values of 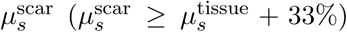, the photon emission profile is so altered that Φ_post_ > Φ_prox_. In all other cases, where Φ_peak_ = Φ_prox_, the reduced decay of radial photon emission is observable as an increase the the FWHM of the emission profile. Changes in 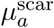 are less observable than 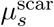 for smaller radii, with a muted effect on Φ_post_, but become more noticeable for larger radii: compare the difference in Φ for *r* ≥ 4 mm, wherein there is negligible difference between all values of 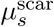 but potentially notable differences for 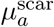.

With the detectability of tissue boundaries established, it is necessary to determine, using realistic parameters, how accurately the location of scar can be established by observed changes in the surface emission profile. Simulations were performed using both semi-infinite tissue, and 8 mm thick tissue (this value being appropriate for left ventricular wall [29]). For all simulations, the scar was simulated to extend from the specified tissue depth to the bottom of the tissue. Scar properties were identical to tissue properties, save for a reduction of *µ*_*s*_ by 50%, which is within the bounds established from the literature—no change in refractive index ensures that no specular reflection occurs.

Figs. 9 and 10 show the surface emission profiles for tissue simulations for both semi-infinite and 8 mm thick tissue, for scars starting at various depths from the tissue surface; fig. 9 demonstrates this for ‘average’ parameters, whereas fig. 10A/10B are for Walton parameters and Fig. 10C/10D are for Ding parameters. Despite the deeply penetrating nature of light for longer wavelengths, there is no noticeable difference between semi-infinite or 8 mm thick tissue for any of the simulations.

**Figure 9:**
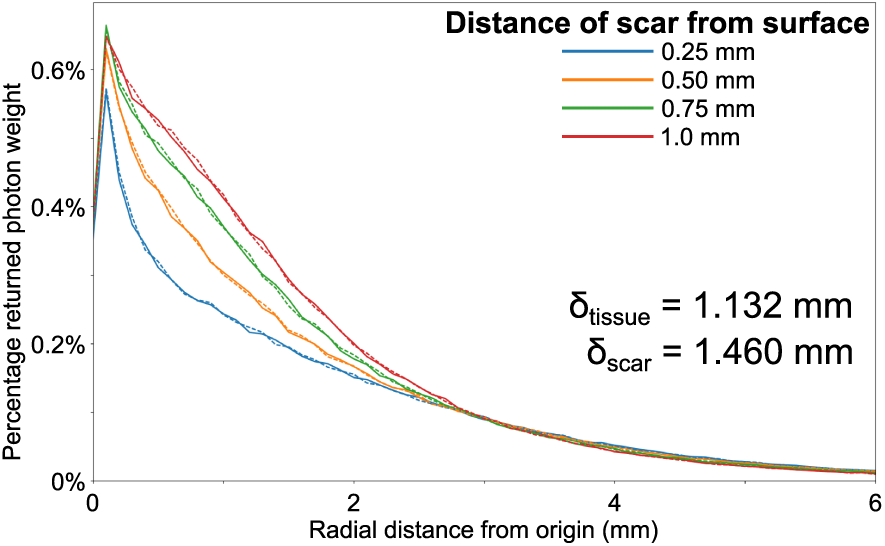
Effect of changing scar depth on surface emission profiles. Effect of scar depth for average optical parameters, with 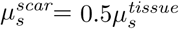. Data shown for both semi-infinite tissue (solid lines) and 8 mm thick tissue (dashed lines). δ values are analytic.

**Figure 10:**
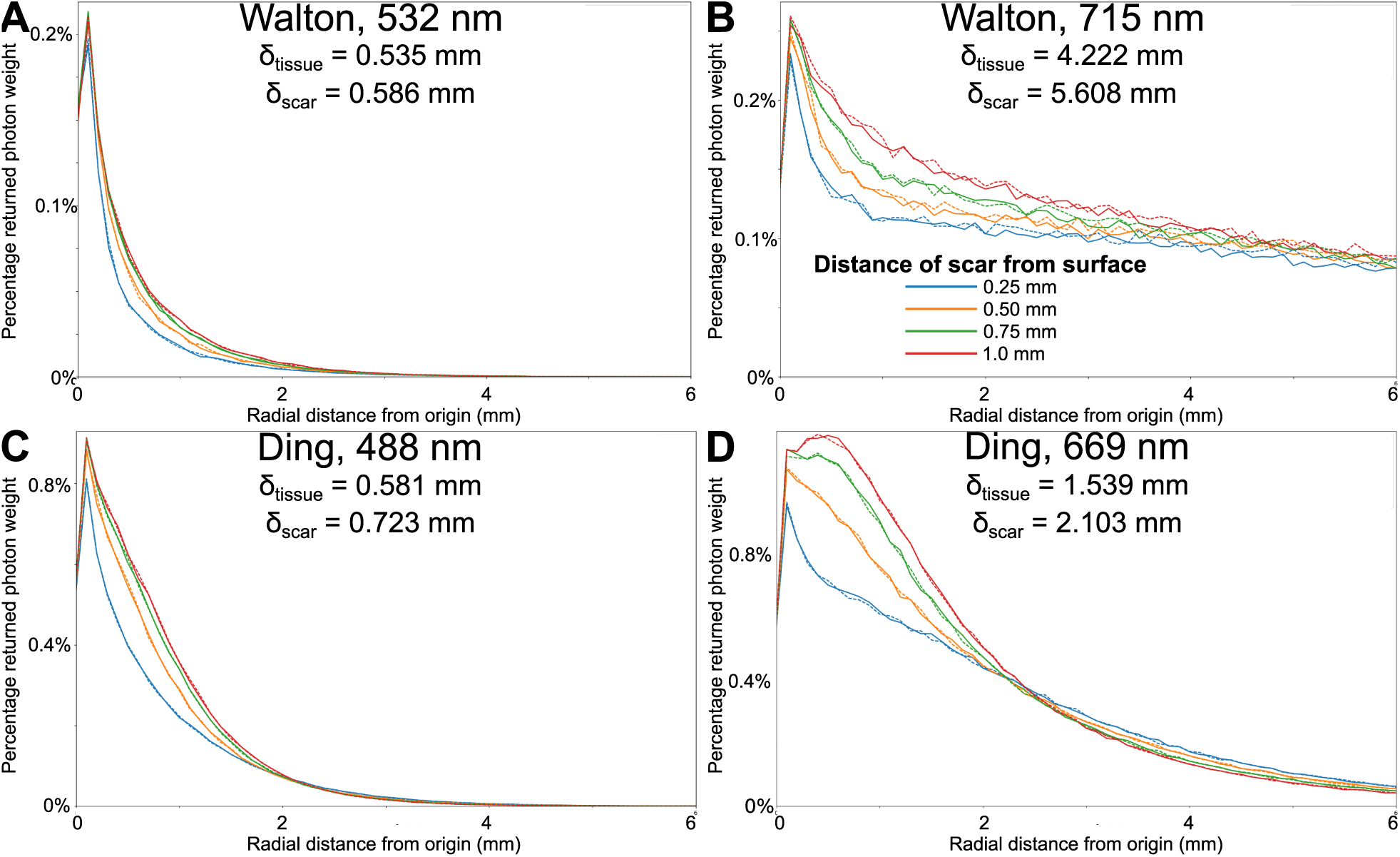
Effect of changing scar properties on surface emission profiles. Effect of scar depth on surface emission profiles, with 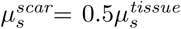, simulated using parameters from [27] (A: 532 nm, B: 715 nm) and [26] (C: 488 nm, D: 669 nm). Data shown for both semi-infinite tissue (solid lines) and 8 mm thick tissue (dashed lines). δ values are analytic.

Under all conditions, as the scar retreats from the surface, Φ_prox_ increases, though the extent to which this is true depends on the parameter choice. With ‘average’ properties, Φ_prox_ increases from 0.57% for a scar 0.25 mm from the surface, to 0.66% for a scar 1 mm from the surface (Fig. 9) (a 16% increase). Both Walton and Ding parameters show ∼ 11% proportional increases, save for 669 nm Ding parameter demonstrating an increase of 16%. Both Walton and Ding simulations indicate that the impact of scar depth on Φ is generally more significant for longer wavelengths. This can be explained by the greater effective penetration depth for those wavelengths.

There are notable differences between the changes in the emission profiles between Walton and Ding parameters. The key difference is in the rate of decay of Φ(*r*) from Φ_prox_ for Walton parameters, which is more rapid when the scar is closer to the surface (it should be noted that this can be observed for 532 nm Walton parameters, but it is far more subtle than at 715 nm). This change in rate of decay can be quantified as a decrease in the FWHM of the surface emission profile.

For the Ding parameters, at 488 nm the rate of decay again increases as the scar moves closer to the surface. However, the profile is more substantially altered for 669 nm, wherein Φ_post_ is more noticeably increased. This increase in Φ_post_ is such that when the scar is 1 mm away from the tissue surface, Φ_post_ > Φ_prox_. This change again results in an increase in the emission profile FWHM. The decay profile for the Ding parameters is such that, for *r* ≳ 2 mm, the emission for simulations when scar is close to the surface is greater than when the scar is distant. It is possible that this inversion is unique to the presence of scar in tissue, but it is unclear whether this difference in Φ would be detectable, as the magnitudes of the emission at these radii is marginal.

## 4 Discussion

This work presents a comprehensive computational study into the feasibility of using optical tomography to establish the structure of cardiac tissue, both in terms of tissue thickness and the presence, and location, of scar. As part of this, an extensive literature search was conducted to establish the experimental ranges for the optical properties of cardiac tissue, and simulations conducted to evaluate the effects of these optical properties. Subsequent simulations assessed the changes to the detectable surface emission profile caused by changes in tissue thickness, and by the presence of scar. The following conclusions can be drawn: (1) it is possible to detect changes in tissue thickness up to ∼ 2 mm; (2) the presence of scar can be detected to similar depths. These effective distances are commensurate with reported effective probe depths for laminar optical tomography [16], [17].

### 4.1 Accurate detection of optical properties

Early simulations were conducted to assess the sensitivity of the surface emission profile to tissue optical parameters, which demonstrated that changes in bulk optical properties can be detected via changes in the surface emission profile. Further, the changes to the profile are subtly different depending on which property varies: changes in scattering dominate the emission profile for small radii from the light source, whereas absorption effects play a more substantial role for larger radii.

### 4.2 Detection of tissue properties

Further simulations demonstrated that changes in the optical properties within the tissue, wherein *µ*_*a,s*_ changes at a given boundary, produce noticeable effects on the emission profile. These changes are due solely to changes in absorption and scattering properties—no change in refractive index was modelled, and thus no specular reflectance occurred. As such, the discontinuity of tissue absorption, and its attendant effect on the surface emission profile, is due entirely to the change in photon path at the boundary, and how the changes in absorption and scattering interact. Depending on the change in individual parameters, the effect of this discontinuity can be striking. The simulations presented here included simulating a discontinuity wherein *µ*_*a,s*_ increase at the tissue boundary. This situation is not expected to reflect the reality for scar: save for one observed increase in 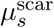 for light wavelengths of 956 nm [20], both parameters are observed to decrease in scar in the literature. However, scar is hardly the only tissue heterogeneity that can cause changes in optical properties: the simulations are agnostic to the physiological cause of the change in optical properties. This can include changes due to the presence of adipose, which is noted to have a substantial effect on optical properties [22], [28], [30], [31], and which can be a substantial component of scar [32].

The results demonstrate noticeable changes in the surface emission profile for both tissue thickness and scar location (with 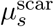 assumed to be half the value for 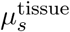). Despite significant penetration of photons into the tissue itself, detectable changes in the profiles are only evident for tissue thickness and scar depths with ∼ 2 mm; this conclusion remains true for more deeply penetrating longer wavelengths. This indicates that this procedure may be more suited to atrial procedures, wherein the effective range of this imaging technique is within the smaller dimensions of the atria, rather than the thicker walls of the ventricles.

It must be noted at the outset that the above conclusions are relevant for all tested optical parameters, and as such they may be assumed to be relatively solid. This is important, as the reported values in literature for optical parameters vary widely, and that the broad conclusions are independent of the specific parameters used lends them credibility. However, the *details* of the changes are sensitive to the parameter choices, which emphasises the need for further study to establish the finer details of any future utility of optical tomography in this setting. While the effect of tissue thickness and presence of scar are less significant than has previously been reported [19], this work was more rigorous in its assessment of optical parameters.

### 4.3 Comparison to alternative methodologies

The clinical relevance of this work is as a method for providing high resolution, real-time measurement of cardiac structure to guide catheter ablation. As such, the results in this work should be compared to alternative methodologies for imaging cardiac structure. One of the extant clinical methodologies is to measure changes in the electrogram of the affected tissue, taking advantage of documented changes in electrical conductivity of scar tissue [1]. However, this suffers from an imprecise mapping between imaging data and catheter position due to cardiac motion, coupled with an inability to capture fine anatomical details. It has also been noted that the presence of scar-related adipose may have significant effects on electrophysiological properties (potentially more so than collagen), in a similar manner to its potential influence here—such influence is currently poorly modelled and understood [33]–[36]. Ultrasound has also been proposed as a promising realtime imaging modality, but suffers from a reduced resolution compared to existing technologies, thus leading to potential inaccuracies in defining lesion boundaries [37].

### 4.4 Limitations and further work

These simulations were conducted in highly simplified models of both myocardium and scar: optical properties were assumed to be homogeneous throughout each type of tissue, and boundaries between the media were well-defined and geometrically simple. Future work would be profitable in using more realistic and complex tissue geometries, both in terms of tissue surface and structure but also in terms of the boundary between tissue and scar. As noted earlier, this work modelled scar as consisting of collagen with a reduced *µ*_*s*_ compared to normal tissue. However, not only can scar possess a far more multi-faceted composition, but scar is not necessarily dense, but can rather exist diffusely, with surviving ‘normal’ myocardium co-existing amongst the scar [38]–[40].

Furthermore, this work is based on an idealised collection geometry, providing complete recovery of the re-emitted surface profile. However, such comprehensive light collection is not realistic, and thus future work could be directed to establish an ideal geometry for a collecting apparatus. However, it can be noted that preliminary analysis conducted during this study indicates that a collection resolution of 1 mm is sufficient to reveal the differences noted in this work.

## 5 Conclusion

The results presented here demonstrate that the surface emission profile for diffusely reflected light is sensitive to the optical parameters within tissue, and furthermore is sensitive to changes of these parameters within the tissue. The changes in the profiles are different depending on which optical parameters change, and dependent on the light wavelength used. Consequently, tissue thickness and presence of scar can noticeably alter surface emission profiles for diffusely reflected light up to depths of ∼ 2 mm. This offers a promising candidate for high resolution, real-time imaging of cardiac structure to guide catheter ablation.

## 6 Supporting Information

A wide range of values for optical parameters are reported in the literature [41]. A summary of these values are presented in S1 Table, S2 Table, S3 Table, S4 Table, with some of the statistics for those values derived from the experimental literature given in S5 Table.

**S1 Table. Literature Values for** *µ*_*a*_ **(mm**^**-1**^). Values obtained from the published literature for the absorption coefficient of cardiac tissue.

**S2 Table. Literature Values for** *µ*_*s*_ **(mm**^**-1**^). Values obtained from the published literature for the scattering coefficient of cardiac tissue.

**S3 Table. Literature Values for** *g*. Values obtained from the published literature for the tissue scattering coefficient of cardiac tissue.

**S4 Table. Literature Values for** *n*_*m*_. Values obtained from the published literature for the refractive index of cardiac tissue.

**S5 Table. Summary of Optical Parameters**. Summary of Optical Parameters Reported in the Literature for Experimental Data From Heart Tissue.

## Notes

* This work was supported by the National Institute for Health Research Biomedical Research Centre at Guy’s and St. Thomas’ National Health Foundation Trust and King’s College, in addition National to the Centre of Excellence in Medical Engineering funded by the Wellcome Trust and Engineering and Physical Sciences Research Council (EPSRC; WT 088641/Z/09/Z). The views expressed are those of the author(s) and not necessarily those of the National Health Service, the National Institute for Health Research, or the Department of Health. The authors acknowledge King’s Health Partners Research and Development Challenge Fund for financial support of this project.

### Competing Interest Statement

The authors have declared no competing interest.

