## Supplemental Information for "Determining cardiac structure by diffuse reflectance of different wavelengths"

### Towards determining cardiac structure by diffuse reflectance of different wavelengths: Supporting Information

July 18, 2018

#### Range of possible model parameters

A wide range of values for optical parameters are reported in the literature [1]. A summary of these values are presented in Tables S1, S2, S3, S4, with some of the statistics for those values derived from the experimental literature given in Table S5.

**Table S1.** Literature Values for $\mu_a$ ($mm^{-1}$).

| Source | Species & Location | Wavelength (nm) | Value | Notes |
|---|---|---|---|---|
| [2] | Human artery (normal) | 514.5 | $1.110 \pm 0.270$ | |
| | | 633 | $0.180 \pm 0.090$ | |
| | | 1060 | $0.090 \pm 0.090$ | |
| | Human artery (plaque) | 514.5 | 1.800 | |
| | | 633 | 0.200 | |
| | | 1060 | 0.140 | |
| [3] | Human artery | 476 | 0.600 | Not experimental. |
| [4] | Endocardium | 1060 | 0.007 | |
| | Epicardium | | 0.035 | |
| [5] | Porcine myocardium (normal) | 1064 | $0.044 \pm 0.005$ | |
| | Porcine myocardium (photocoagulated) | | $0.051 \pm 0.009$ | |
| [6] | Canine myocardium (normal) | 1064 | $0.040 \pm 0.020$ | |
| | Canine myocardium (photocoagulated) | | $0.035 \pm 0.016$ | |
| | Human myocardium (normal) | | $0.030 \pm 0.020$ | |
| | Human epicardium (fat) | | $0.021 \pm 0.016$ | |
| | Human myocardium (scar) | | 0.040 | |
| [7] | Canine myocardium | 632.8 | $0.200 \pm 0.040$ | |
| [8] | Canine myocardium | 630 | $0.200 \pm 0.020$ | |
| | | 632.8 | $0.210 \pm 0.010$ | |
| | | 790.0 | $0.098 \pm 0.020$ | |
| [9] | Sheep heart | 670 | 0.478 | Estimated using iterative Monte Carlo modelling to match experimental data. |
| [10] | Human cerebellum (normal) | 674 | $0.250 \pm 0.050$ | |
| | | 849 | $0.095 \pm 0.020$ | |
| | | 956 | $0.090 \pm 0.020$ | |
| | Human cerebellum (scar) | 674 | $< 0.02$ | |
| | | 849 | $< 0.02$ | |
| | | 956 | $< 0.02$ | |
| [11] | Rabbit heart | 488 | $0.710 \pm 0.240$ | Using Rh237 & Oregon dye. |
| | | | $0.520 \pm 0.130$ | Using Di-4-ANEPPS dye. |
| | | 669 | $0.150 \pm 0.100$ | Using Rh237 & Oregon dye. |
| | | | $0.100 \pm 0.040$ | Using Di-4-ANEPPS dye. |
| [12] | Goat heart | 670 | 0.127 | Estimated using iterative Monte Carlo modelling to match experimental data. Also used in [13]. |
| | Goat adipose | | 41.992 | |
| [14] | Porcine myocardium (normal) | 633 | 0.200 | |
| | Porcine myocardium (photocoagulated) | | 0.270 | |
| | Porcine myocardium (normal) | 790 | 0.062 | |
| [15] | Rat myocardium | 610 | 0.165 | Also used in [16]. |
| | | 532 | 0.880 | Also used in [16, 17]. |
| [16] | Rat | 660 | 0.070 | Not experimental, also used in [17]. |
| | | 715 | 0.050 | |
| [18] | Rat heart | 710 | $0.270 \pm 0.060$ | $\mu_a$ calculated from entire bandwidth. |

**Table S2.** Literature Values for $\mu_s$ ($mm^{-1}$).

| Source | Species & Location | Wavelength (nm) | Value | Notes |
|---|---|---|---|---|
| [2] | Human artery (normal) | 514.5 | $1.10 \pm 0.08$ | |
| | | 633 | $0.63 \pm 0.14$ | |
| | | 1060 | $0.28 \pm 0.02$ | |
| | Human artery (plaque) | 514.5 | 1.90 | |
| | | 633 | 1.20 | |
| | | 1060 | 0.23 | |
| [3] | Human artery | 476 | 41.40 | Not experimental. |
| [4] | Endocardium | 1060 | 13.60 | |
| | Epicardium | | 16.70 | |
| [5] | Porcine myocardium (normal) | 1064 | $0.427 \pm 0.033$ | |
| | Porcine myocardium (photocoagulated) | | $1.743 \pm 0.101$ | |
| [6] | Canine myocardium (normal) | 1064 | $17.35 \pm 2.21$ | |
| | Canine myocardium (photocoagulated) | | $24.59 \pm 2.33$ | |
| | Human myocardium (normal) | | $17.75 \pm 3.75$ | |
| | Human epicardium (fat) | | $12.71 \pm 1.86$ | |
| | Human myocardium (scar) | | 13.7 | |
| [7] | Canine myocardium | 632.8 | $16.00 \pm 3.00$ | |
| [8] | Canine myocardium | 630 | $15.90 \pm 0.60$ | |
| | | 632.8 | $19.10 \pm 1.60$ | |
| | | 790.0 | $16.40 \pm 1.00$ | |
| [9] | Sheep heart | 670 | 5.54 | Estimated using iterative Monte Carlo modelling to match experimental data. |
| [10] | Human cerebellum (normal) | 674 | $27.00 \pm 2.00$ | Estimated using $g = 0.95$. |
| | | 849 | $17.00 \pm 2.00$ | |
| | | 956 | $15.60 \pm 2.00$ | |
| | Human cerebellum (scar) | 674 | $13.00 \pm 1.00$ | |
| | | 849 | $16.00 \pm 1.00$ | |
| | | 956 | $13.00 \pm 1.00$ | |
| [11] | Rabbit heart | 488 | $29.20 \pm 4.90$ | Using Rh237 & Oregon dye. |
| | | | $23.00 \pm 2.50$ | Using Di-4-ANEPPS dye. |
| | | 669 | $27.10 \pm 4.00$ | Using Rh237 & Oregon dye. |
| | | | $21.80 \pm 2.50$ | Using Di-4-ANEPPS dye. |
| [12] | Goat heart | 670 | 10.03 | Estimated using iterative Monte Carlo modelling to match experimental data. Also used in [13]. |
| | Goat adipose | | 0.15 | |
| [14] | Porcine myocardium (normal) | 633 | 10.00 | |
| | Porcine myocardium (photocoagulated) | | 15.00 | |
| | Porcine myocardium (normal) | 790 | 10.30 | |
| [15] | Rat myocardium | 610 | 6.30 | Also used in [16]. |
| | | 532 | 7.42 | Also used in [16,17]. |
| [16] | Rat | 660 | 5.90 | Estimated from reduced $mu_s$ using $g = 0.9$. |
| | | 715 | 5.40 | |
| [18] | Rat heart | 710 | $13.26 \pm 0.37$ | $\mu_s$ calculated from entire bandwidth. |
| [17] | Myocardium | 532 | 14.80 | Based on [16], with altered $g$. |
| | | 660 | 11.80 | |
| | | 745 | 10.80 | |

**Table S3.** Literature Values for $g$.

| Source | Species & Location | Wavelength (nm) | Value | Notes |
|---|---|---|---|---|
| [3] | Human artery | 476 | 0.910 | Not experimental. |
| [4] | Endocardium | 1060 | 0.973 | Reported second hand. |
| | Epicardium | | 0.983 | |
| [6] | Canine myocardium (normal) | 1064 | $0.974 \pm 0.008$ | |
| | Canine myocardium (photocoagulated) | | $0.956 \pm 0.016$ | |
| | Human myocardium (normal) | | $0.964 \pm 0.005$ | |
| | Human epicardium (fat) | | $0.933 \pm 0.017$ | |
| | Human myocardium (scar) | | 0.975 | |
| [7] | Canine myocardium | 632.8 | $0.930 \pm 0.020$ | |
| [8] | Canine myocardium | 630 | $0.854 \pm 0.015$ | |
| | | 632.8 | $0.940 \pm 0.004$ | |
| | | 790.0 | $0.943 \pm 0.004$ | |
| [11] | Rabbit heart | 488 | $0.930 \pm 0.020$ | Using Rh237 & Oregon dye. |
| | | | $0.940 \pm 0.020$ | Using Di-4-ANEPPS dye. |
| | | 669 | $0.950 \pm 0.010$ | Using Rh237 & Oregon dye. |
| | | | $0.960 \pm 0.010$ | Using Di-4-ANEPPS dye. |
| [12] | Goat heart | 670 | 0.995 | Estimated using iterative Monte Carlo modelling to match experimental data. |
| | Goat adipose | | 0.994 | Estimated using iterative Monte Carlo modelling to match experimental data. Also used in [13]. |
| [14] | Porcine myocardium | 633 | 0.900 | Not experimental. |
| | Porcine myocardium | 790 | 0.930 | |
| [15] | Rat myocardium | 610 | 0.900 | |
| | | 532 | 0.900 | |
| [18] | Rat heart | 710 | $0.960 \pm 0.020$ | |
| [17] | Myocardium | 532<br>660<br>745 | 0.950 | Not experimental. |
| [13] | Goat heart | 632.8 | 0.990 | Not experimental. |

**Table S4.** Literature Values for $n_m$.

| Source | Species & Location | Wavelength (nm) | Value |
|---|---|---|---|
| [19] | Bovine skeletal muscle | 390 | 1.425 |
| | | 700 | 1.388 |
| | | 632.8 | $1.412 \pm 0.006$ |
| | Bovine adipose | | 1.455 |
| [3][1] | Human artery | 476 | 1.370 |
| [18][1] | Rat heart | 710 | 1.380 |
| [17][1] | Myocardium | 532 | 1.400 |
| | | 660 | |
| | | 745 | |
| [13][1] | Goat heart (normal) | 632.8 | 1.400 |
| | Goat heart (fat) | | 1.455 |

[1] Not experimental

**Table S5.** Summary of Optical Parameters Reported in the Literature for Experimental Data From Heart Tissue

| Parameter | $n$ | Mean Value | Min. Value | Max. Value |
|---|---|---|---|---|
| $\mu_a$ (mm$^{-1}$) | 23 | 0.1886 | 0.007 | 0.88 |
| $\mu_s$ (mm$^{-1}$) | 23 | 15.1891 | 0.427 | 29.2 |
| $g$ | 19 | 0.9418 | 0.854 | 0.983 |

$n$ is the number of values from the literature used to generate the data.
